# Oocyte exposure to low levels of triclosan has a significant impact on subsequent embryo physiology

**DOI:** 10.1101/2025.05.01.651623

**Authors:** Vasiliki Papachristofi, Paul McKeegan, Henry J Leese, Jeanette M. Rotchell, Roger Sturmey

**Affiliations:** University of Hull; University of Lincoln

## Abstract

Triclosan (TCS) is an anti-microbial agent in a wide range of health care products. It has been found in various human bodily fluids and is a potential reproductive toxicant. However, the effect of TCS on early embryo development in mammalian species is limited. We therefore asked whether exposure to TCS affects mammalian cumulus-oocyte-complexes (COCs), and if so, whether the effects persist into the early embryo. COCs, isolated from abattoir-derived bovine ovaries were exposed to two environmentally relevant doses of TCS (1 nM and 10 nM) during in vitrao maturation. When exposed to 1 nM TCS during in vitro maturation, progesterone release from bovine oocytes was elevated. Furthermore, altered pyruvate metabolism and mitochondrial dysfunction also observed; specifically, O_2_ consumption coupled to ATP production was significantly decreased in COCs after acute exposure of TCS prior to maturation, whereas proton leak from the respiratory chain was increased. Subsequently, TCS-exposed COCs were fertilised. Fewer oocytes were able to develop to blastocyst when exposed to 1 nM TCS during maturation compared to the Control group (12.22±% vs 29.11±% n=5; *p*=0.02) and those that did reach the blastocyst displayed impaired glycolytic and amino acid metabolic activity. These findings indicate for the first time that oocytes exposed to TCS during the final stages of maturation give rise to embryos with impaired mitochondria function, altered steroidogenesis and disrupted metabolic activity.

## Introduction

Triclosan (TCS) is a broad-spectrum agent with antimicrobial activity, commonly found in products including soaps, healthcare antiseptic products such as mouthwash, toothpaste, personal hygiene products including acne cream, skin cream, sunscreens and deodorant, and medical products including surgical sutures, catheters and ureteral stents (Rodricks *et al*., 2010; Dann and Hontela, 2011; Ye *et al*., 2011). Although the use of TCS was restricted in specific certain wash products in the USA in 2016 (FDA, 2016) and from products used in hospital in 2017 (FDA, 2017), inclusion of TCS remains permitted in personal care products, cosmetics, clothes, and toys. Because of this, the daily TCS exposure of the general population is still estimated to be high, with geometric mean from 0.7 μg/L to 7.85 μg/L depending on the country (Iyer *et al*., 2018). As a consequence of extensive inclusion in consumer products, TCS has been widely reported to have accumulated in the aquatic and terrestrial environment (Chalew and Halden, 2009; Fair *et al*., 2009; Houten and Wanders, 2010). Such bioaccumulation in aquatic biota has prompted concerns about the impact of TCS on human and animal health (Olaniyan *et al*., 2016).

Of particular concern, TCS has been reported to be present in a range of biological fluids. For example, TCS was detected at nanomolar concentrations in over 2000 urine samples collected during a study of the USA general population (Calafat *et al*., 2008). Similar reports have confirmed the presence of TCS and its conjugated forms in the urine of population cohorts from India (Xue *et al*., 2015), Spain (Azzouz *et al*., 2016) and Belgium (Pirard *et al*., 2012). Furthermore, in a study of 1,890 first-trimester urine samples of pregnant women, free TCS and TCS-metabolites (with geometric means 0.5μg/L and 12.30μg/L, respectively) were detected in more than 75% of samples examined (Arbuckle *et al*., 2015). Perhaps most startlingly, TCS has been identified in blood (4.1–41.4 nM;(Azzouz *et al*., 2016)), serum (0.035–1200 nM; (Allmyr *et al*., 2008)) and breast milk over a wide concentration range (0.062–252 nM) (Adolfsson-Erici *et al*., 2002; Dayan, 2007; Azzouz *et al*., 2016; Bever *et al*., 2018)). Additionally, TCS has been detected in human tissues, including adipose tissue, brain and liver at levels of 0.23 ng/g −29.03 ng/g (Geens *et al*., 2012) and a recent study reported the presence of TCS in the ovarian follicle at a concentration of ∼100pg/ml (Li et al 2024). Of critical importance, there are reports of correlations between increased urinary concentration of TCS (8.9 – 19.1 μg/L) and diminished ovarian reserve (Mínguez-Alarcón *et al*., 2017; Jurewicz *et al*., 2019) and of follicular TCS levels and embryo quality (Li et al 2024), hinting at a link between TCS and fertility.

Beyond studies of TCS accumulation uptake, the biological implications of such exposure are now becoming apparent. For instance, TCS exposure has been correlated with oxidative stress and mitochondrial dysfunction in porcine oocytes (Park *et al*., 2020; Zhao *et al*., 2025) and zebrafish embryos (Fu *et al*., 2021; Liu *et al*., 2022). Indeed, a study in human epithelial cell lines showed that TCS (1.25 −50 μM) could impair mitochondria function by stimulating superoxidase release and inhibiting complex II (Teplova *et al*., 2017) suggesting a direct effect of TCS on cellular metabolic processes. Furthermore, TCS exposure in the micromolar (μΜ) range seems to be correlated with impaired function of early reproductive events such as reduced meiotic maturation in pig oocytes (Zhao *et al*., 2025), and in the implantation process through an effect on uterine receptivity (Dong *et al*., 2022) and on preimplantation embryo development by reducing the expression of pluripotent markers (Yang *et al*., 2022) in mice.

Oocyte quality is a major determinant of embryo developmental competence. Disruption of COC physiological function can have subsequent effects on the physiology and metabolic function of subsequent early embryos as reviewed by Krisher, (2004); and Keefe *et al*., (2015). This is important because impaired metabolic function in embryos has been correlated with decreased developmental outcomes in mouse (Gardner and Leese, 1987; Houghton *et al*., 1996), bovine (Guerif *et al*., 2013) and human, embryos (Houghton *et al*., 2002; Brison *et al*., 2004). Nutritional markers, such as glucose and pyruvate depletion and lactate production, and animo acid turnover (Martin, 2000; Leese, 2003), as well as physiological markers such as oxygen consumption rate (Lopes *et al*., 2007; Ottosen *et al*., 2007) have previously been associated with embryo quality and developmental potential. Underscoring this, recent evidence ties appropriate metabolic function to genome activation (Nagaraj *et al*., 2017), illustrating the fundamental importance of metabolism to early development. Beyond this, metabolic activity of embryos may have links to the lifelong health of the resulting offspring. The Developmental Origins of Health and Disease (DOHaD) hypothesis (Barker, 2007) has proposed that certain environmental stimuli (such as nutrition, stress, chemical exposure) during essential/key? developmental periods may induce phenotypic adaptation during the preimplantation stages of development that persist into later life (Watkins *et al*., 2008, 2010; Eckert *et al*., 2012).

Given the intrinsic importance of gamete and embryo metabolism, the aim of this study was to examine the extent to which steroidogenesis and metabolic activity of bovine COCs during In Vitro Maturation (IVM) are sensitive to the reproductive toxicant TCS. In addition, the embryo developmental potential and the metabolic profile of the resulting embryos was assessed at the blastocyst stage. The bovine model was chosen since it has been identified as an effective model for reproductive toxicological studies due to its functional and physiological similarities with the humans (Ménézo and Hérubel, 2002; Santos *et al*., 2014). Importantly, this study has focussed on physiologically-relevant doses of TCS.

## Material and Methods

All work was carried out after ethical review from the Institutional Ethical Review Board (Ref HYMS 18 47).

### Experimental design overview

COCs, isolated from abattoir-derived ovaries, were cultured in the presence of two different doses of TCS (10nM and 1nM) during in vitro maturation. Both doses are within the range previously detected in human bodily fluids. The levels of steroid hormones (17β-estradiol and progesterone) and key-metabolites (pyruvate, lactate) were quantified in spent media using ELISA and microfluorometric assays, respectively. Furthermore, using extracellular flux analysis, the Oxygen Consumption Rate (OCR) was measured as an indicator of the mitochondrial function of COCs (Muller et al 2019) acutely exposed to TCS <u>prior to</u> <u>maturation.</u>

Subsequently, the TCS-exposed COCs were fertlilised and cultured for in vitro embryo production without further TCS addition. The rates of embryo cleavage and blastocyst formation were recorded on Day 2 and Day 7/Day 8, respectively and the metabolic activity of the resulting blastocysts determined. The key energy metabolites glucose, pyruvate, lactate, amino acids turnover were quantified in spent media using microfluorometric assay and HPLC respectively.

### Cumulus oocyte complexes’ in vitro maturation (IVM) and in vitro culture of bovine embryos (IVC)

Bovine reproductive tracts were collected from a local abattoir and transferred to the laboratory within 2 hours from the animals’ slaughter. Upon arrival i the laboratory, the ovaries were isolated from the rest of the reproductive tract and wnashed three times in pre-warmed PBS supplemented with Antimycotic-Antibiotic at 39°C. IVM and IVC were performed as described previously (Orsi and Leese, 2004). Briefly, follicles with a diameter of between 2-8mm were punctured and the contents aspirated into M199 media supplemented with HEPES buffer, heparin and bovine serum albumin. Oocytes with at least 2 layers of intact cumulus cells were selected, washed and placed in groups of 35 – 45 into 500 μl bovine maturation media, which was prepared in house from M199 media supplemented with NaHCO[, gonadotrophins FSH and LH and growth factors EGF and FGF. COCs were incubated in a humidified environment, in 5% CO_2_, in air for 21 to 23 hours. After this time, mature COCs were used for in vitro fertilisation (IVF) and the spent maturation medium was stored at −80°C until further analysis, described in the following sections.

For IVF, semen samples stored in liquid nitrogen, from a single bull with proven fertility, were thawed instantly in a warm bath at 39°C. The semen samples were transferred in Semen-Prep commercial media (IVF Bioscience) and centrifuged twice at 328 x *g* for 5 min. Groups of 35-45 oocytes were co-incubated with 1×10^6^ sperm cells in fertilisation-TALP media for 18-22 hours in 5% CO_2_, in air.

Putative zygotes were selected and vortexed vigorously for 2 min to remove any remaining cumulus cells before being placed in groups of 20 to 25 in 30 μl of Synthetic Oviduct Fluid media supplemented with amino acids and bovine serum albumin, BSA (SOFaaBSA) under mineral oil and incubated for maximum of 8 days under hypoxic conditions (5% CO_2_, 5% O_2_ Bal N_2_). On day 2 and day 7/day 8 cleavage and blastocyst rates of the embryos were recorded respectively. On day 7 and day 8, the blastocysts were retrieved for individual embryo culture described in the following section.

### Preparation and storage of TCS stocks

For the purposes of all experiments -except for those of Oxygen Consumption Rate measurement-TCS (CAS: 3380-34-5) was diluted into 100% DMSO (CAS: 67-68-5) creating a stock of 500x the final concentration and stored in aliquots at −20°C in glass vials for a maximum of 3 months in order to prevent TCS degradation. On the day of IVM the TCS stock was further diluted into maturation medium for a final concentration of 10 nM and 1 nM and incubated in 5% CO_2_, in air at 39°C for at least 2 hours prior to COCs addition. TCS exposure took place solely during oocyte maturation. Where appropriate, vehicle controls were prepared with DMSO added to maturation medium in the absence of TCS.

### Individual in vitro culture of embryos for metabolic assays

On day 7 and day 8 of embryo culture, blastocysts were washed twice and then cultured individually in SOF “analysis” media in 5μl droplets under oil for approximately 22-24 hours in hypoxic conditions of 5% CO_2_ and 5% O_2_. SOF “analysis” medium is an identical version of SOFaaBSA in all aspects except for glucose and lactate concentrations which are 0.5 mM and 0 mM, respectively (Geurif *et al.,* 2013). The morphological stage and precise time of transfer of blastocysts into and out of the SOF analysis medium were recorded. At the conclusion of embryo culture, the blastocysts were removed, and the incubation plates sealed and stored at −80°C until analysis. Finally, COnsuption and RElease analysis (CORE analysis) of key metabolites and amino acids was performed as described in (Guerif *et al*., 2013).

### Determination of steroids hormone release

Spent media from COCs exposed to TCS during maturation were collected and stored at - 80oC for steroid hormone quantification. 17β-estradiol and progesterone concentrations in the spent media were quantified using enzyme immunoassay reagents (Enzo Life Science, Farmingdale, NY, USA,). The reported sensitivity of 17β-estradiol ELISA kit (ADI-900-008) was 28.5 pg/ml and the inter-assay and intra-assay coefficients of variation ranged from 5.2 to 7.4% and 8.4 to 9.2%, respectively. The limit of detection of the progesterone ELISA kit (ADI-900-01) was 8.57pg/ml and the inter-assay and intra-assay coefficients of variation varied from 2.7 to 6.8% and 4.9 to 7.6%, respectively. The quantification of hormone concentrations in the samples was performed according to manufacturer’s instructions and subsequently normalised against the number of COCs and the hours of incubation per spent media sample expressed as pmol/hour/COC.

### Identification of COCs nuclear status after IVM

COCs were collected after IVM, washed three times in 0.2% PVP-PBS and then vortexed vigorously to remove remaining cumulus cells. Finally, the denuded oocytes were labelled with Hoechst-EtOH nuclear staining and incubated in 4°C overnight, after which the nuclear status was assessed using a fluorescence microscope (Axiocam 506, Zeiss).

### Quantification of consumption and release of key metabolites’ (CORE analysis)

The quantification of pyruvate and lactate in the conditioned maturation medium, and glucose, pyruvate, and lactate in spent SOF analysis media were performed using a microfluorometric assay as described previously (Guerif et al., 2013). Briefly, 1μl of sample or blank medium was added into 9 μl of relevant-assays’ mixture and the concentation of the key metabolites were determined indirectly using the enzymatic-induced oxidation/reduction of NAD(P)H, which was measured fluorometrically. The pyruvate assay mixture comprised 0.1 mM NADH and 40 IU/ml lactate dehydrogenase in 4.6 mM EPPS buffer, pH 8.0, which was co-incubated with samples at 37°C for 3 minutes. The final concentrations in each sample were determined against a six-point standard curve from 0–0.45 mM pyruvate for BMM media and 0-0.36mM for SOF analysis media. The lactate mixture contained 40 IU/ml lactate dehydrogenase (LDH) in a glycine-hydrazine buffer, pH 9.4, which was co-incubated with samples at 37°C for 30 minutes. A six-point standard curve of lactate was used to determine the lactate concentation in the spent media. A 0 to 2.5 mM and a 0 to 1.25 mM standard curve was run against BMM samples and SOF analysis samples, respectively. Finally, the glucose mixture included 0.4 mM dithiothreitol, 3.07 mM MgSO_4_, 0.42 mM ATP, 1.25 mM NADP^+^, 20 IU /ml hexokinase/glucose-6-phosphate dehydrogenase (HK/G6PDH) in EPPS buffer at pH 8.0. Samples were co-incubated at 37°C for 10 minutes. The final concentration of glucose in the SOF analysis media was calculated based on a six point, 0 to 0.5mM, standard curve. After the final concentration was calculated further normalisation was performed against the hours of incubation for COCs or blastocyst and the number of COCs within a droplet. Blastocysts were cultured singly. Data are expressed in pmol/hour/COC or pmol/hour/embryo respectively.

### Quantification of amino acids turn over

Reverse Phase- High Performance Liquid Chromatography was used as described previously (Houghton *et al*., 2002) for the quantification of 18 amino acids in the spent SOF analysis media. Briefly, the detection of the amino acid is based on different size and hydrophobicity of the amino acids. The amino acids underwent pre-column derivatisation with O-phthaldialdehyde (OPA), supplemented with 1mg/ml 2-mercaptoethanol, which resulted in the formation of amino acid conjugates with OPA and generated a fluorescent signal, detectable at 450 nm.

Aspartic acid, glutamic acid, asparagine, serine, histidine, glutamine, glycine, threonine, arginine, alanine, tyrosine, tryptophan, methionine, valine, phenylalanine, isoleucine, leucine and lysine were quantified with this technique calculated using the retention time (RT) and the area under the curve (AUC), respectively, and compared to standards. Certified amino acids standards were used (AAS18, Sigma Aldrich, Burlington, MA, United States) supplemented with asparagine, glutamine, tryptophan, and 2-Diethylomino-N-3-phenylmethoxypheny-l-acetamide (D-ABA) which were made in-house. D-ABA was used as internal standard, as a non-metabolisable amino acid. The concentration of the amino acids in the final standard solution was 12.5μΜ. The samples were diluted to 1:12.5 in ddH2O and analysed with a single injection per vial.

The concentration of each amino acid in each sample was further subtracted from the concentration of the same amino acid in the blank drop and finally normalised against the recorded embryo incubation time and the results expressed as pmol/hour/embryo.

### Determination of mitochondria function using Oxygen Consumption Rate

The mitochondrial function after acute TCS exposure of oocytes prior to maturation was determined using the measurement of Oxygen Consumption Rate (OCR) and its components with Extracellular Flux Analysis (EFA) as reported by (Muller *et al*., 2019). Briefly, using a sensor-containing Seahorse fluxpak, OCR and its components were measured in a Seahorse-XFp analyser (Agilent, Santa Clara, California). For each assay, an overnight pre-incubation in a non-CO2 humidifier and an ‘on-the-day calibration’ of the sensor-containing Seahorse fluxpak were included. Upon completion of these steps, COCs were loaded into cell plate and placed into the Seahorse-XFp.

For the purpose of this experiment, a 17-point measurements protocol was developed. Each point of measurement included a three-minute measurement and one-minute wait period. During the 3-minute measurement period, the sensor was lowered creating an airtight microenvironment able to detect the depletion of dissolved oxygen. During the 1-minute waiting step the sensor was raised up allowing re-calibration. During the course of the 17-point protocol, 4 serial injections occurred: 1 TCS and 3 mitochondrial inhibitors (oligomycin, FCCP, and Rotenone/Antimycin A). During the first 3 points no injection took place, TCS was injected after the completion of the 3^rd^ point, Oligomycin after 8^th^ point, FCCP was added after the 11^th^ point and a mixture of Rotenone/Antimycin A injected after 14^th^ point. The final concentration of TCS in the cell wells was 0 nM, 1 nM or 10 nM depending on the group, 1μM for oligomycin 5μM FCCP, and finally 2.5μM of Rotenone/ Antimycin A mixture as suggested by Muller et al., 2019. The results of OCR assay, expressed as pmol/min/well were normalised to oocyte number per well.

### Statistical analysis

All data were normalised against the number of the COCs or embryos and the hours of incubations as described above in each section of the methods. Proportional data underwent Arc-sine transformation prior to any statistical test. The normal distribution of each non-proportional data set was checked using the D’Agostino & Pearson test or Shapiro-Wilk in the cases where the sample size was too small for the D’Agostino & Pearson test. The data sets that failed the normality test were log transformed and parametric tests (one-way ANOVA or two-way ANOVA) were used to identify possible differences. In the cases where log transformation was not possible non-parametric tests were used. One-way ANOVA followed by post-hoc Tukey test or Kruskal-Wallis test followed by Dunn’s test were used when more than two groups were compared. Two-way ANOVA followed by post-hoc Tukey test or multiple Mann-Whitney was used when each experimental group included more than one condition. All statistical tests were performed on GraphPad Prism using a significant difference *p* value < 0.05. All graphs are represented as mean ± SD.

## Results

COCs were first exposed to two different concentrations of TCS during maturation and hormone release prior to the determination of metabolic activity. Release of 17β-estradiol by bovine COCs in vitro was not influenced by the presence of TCS regardless of the administrated dose (Fig 1a). Progesterone production was not influenced by the high dose of TCS (10 nM) however, COCs exposed to 1 nM TCS released significantly more progesterone than non-treated controls (2.85 vs 4.78 pg/hour/oocyte p=0.034; Figure 1).

**Figure 1:**
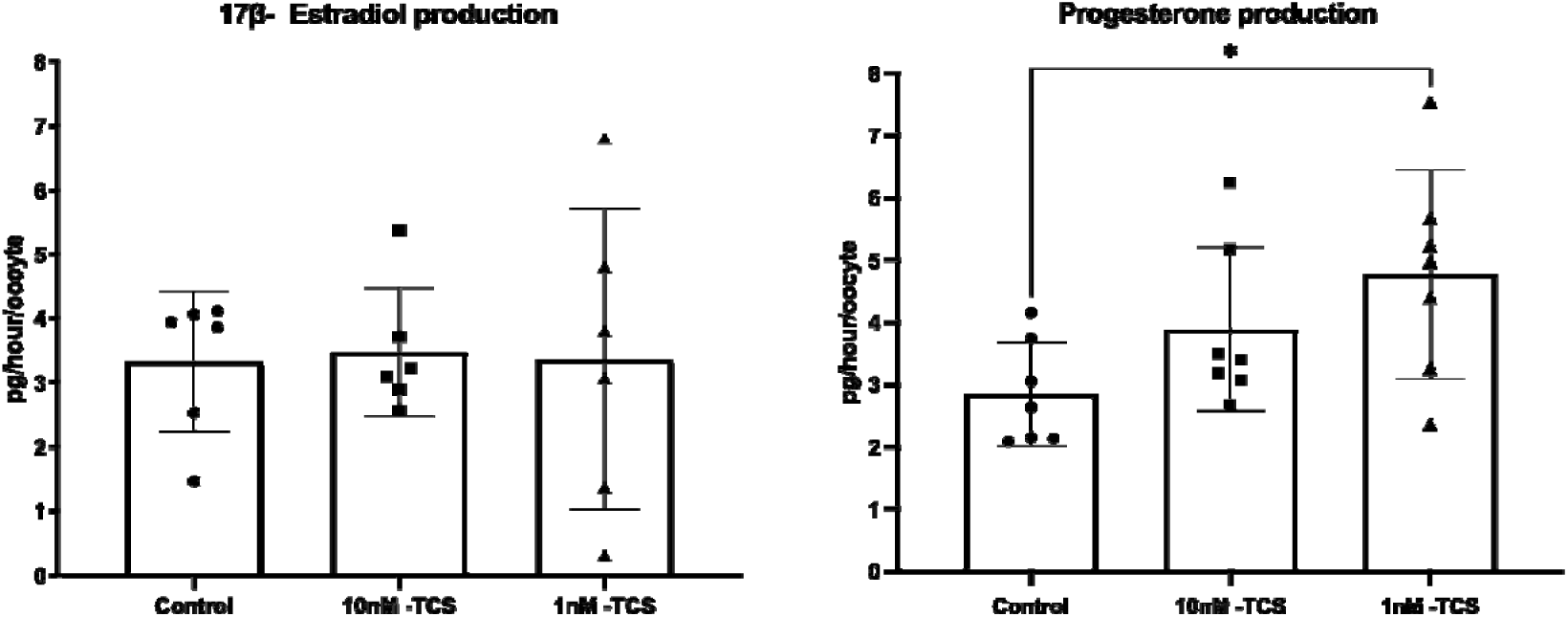
Release of 17β-estradiol and progesterone from bovine COCs exposed to two different concentrations of TCS at 10 nM or 1 nM during in vitro maturation (n=6 each dot represents an individual replicate). Differences were assessed using one-way ANOVA followed by Tukey test. * p<0.05 Data are presented as mean ± SD

Next, the energy substrates, pyruvate production/depletion and lactate production, by COCs during IVM were recorded (Fig 2). The findings revealed that COCs cultured without TCS or with the vehicle solvent (DMSO) released pyruvate (82.84 ±33.9 pmol/hour/COCs and 73.97 ±56.1 pmol/hour/COCs respectively) whilst pyruvate was depleted from culture media by COCs when exposed to TCS p<0.0001 (Figure 2). COCs exposed to 10nM dose consumed an average of 112.8 ±35.8 pmol/hour/COCs pyruvate and those exposed to 1nM exhibited a mean pyruvate consumption of 106.4 ±27.1 pmol/hour/COCs. Notably, TCS had no effect on lactate release (Fig 2).

**Figure 2:**
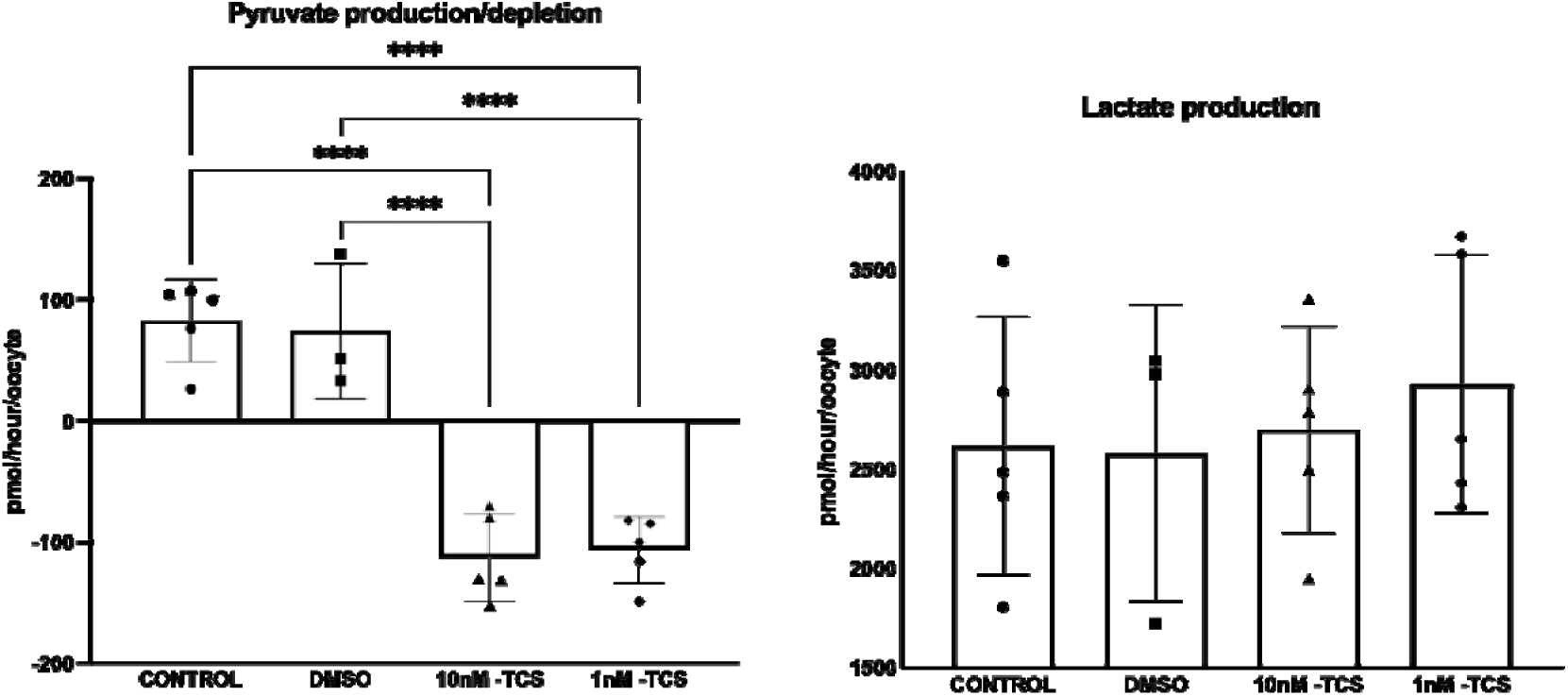
Depletion/production of pyruvate and lactate by bovine COCs after exposure to two concentrations of Triclosan (10nM, 1nM) during in vitro maturation (n=5 individual replicates with between 25-35 oocytes per replicate). Differences were tested for significance using one-way ANOVA followed by Tukey test. * p<0.05; ** p<0.01; *** p<0.001; **** p<0.0001. Data presented in mean ± SD

In addition, the nuclear status of 352 COCs in total after IVM was assessed (Table 1). The values corresponded to 3 biological replicates. TCS exposure did not affect the proportion of COCs reaching MII stage of maturation. (Fig 3).

**Table I:**
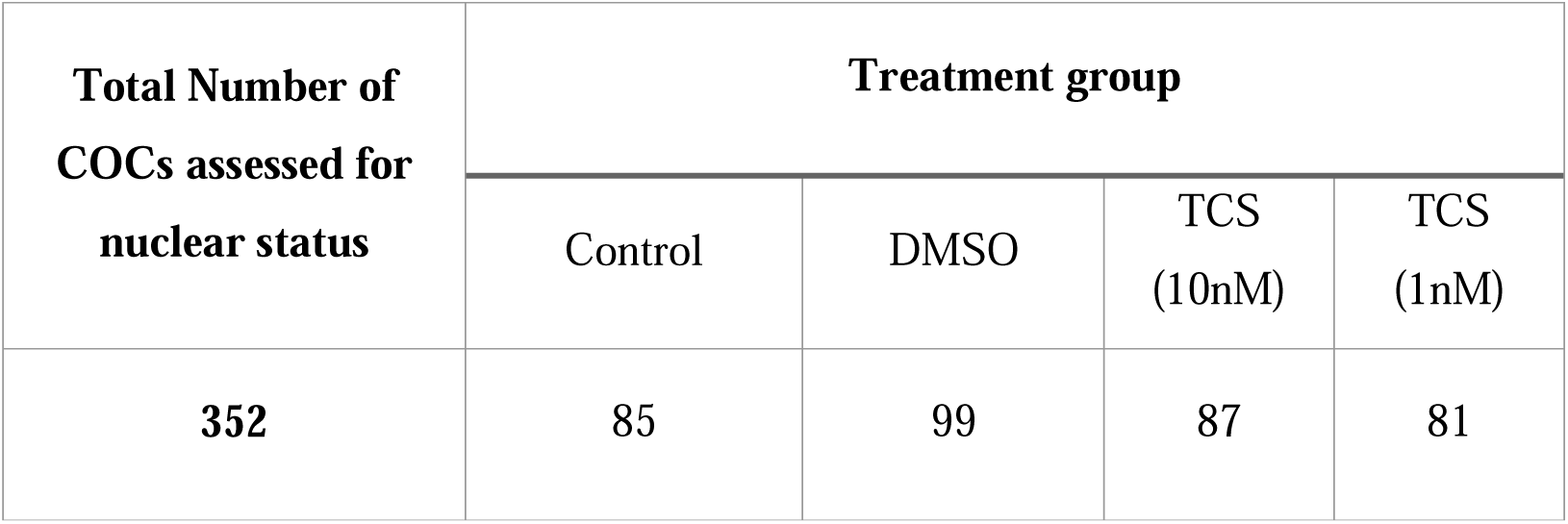
Number of COCs assessed for the nuclear status per group of treatment.

**Table II:** Turnover of each individual amino acid per treatment group. Amino Acids in bold with an asterisk * are those with statistically significant difference. The results are expressed on pmol/hour/embryo ± SD The **amino acids**\* are illustrated in Figure 7

| Amino Acids | Treatment |  |  |  |  |  |  |  |
| --- | --- | --- | --- | --- | --- | --- | --- | --- |
|  | Control |  | DMSO |  | TCS (10nM) |  | TCS (1nM) |  |
|  | Mean | SD | Mean | SD | Mean | SD | Mean | SD |
| Asp | 0.068 | $\pm 0.147$ | 0.113 | $\pm 0.237$ | 0.043 | $\pm 0.273$ | 0.025 | $\pm 0.178$ |
| Glu | -1.031 | $\pm 0.771$ | -1.083 | $\pm 0.727$ | -1.209 | $\pm 0.943$ | -1.32 | $\pm 0.745$ |
| **Asn\*** | **0.035** | **$\pm 0.139$** | **0.093** | **$\pm 0.382$** | **-0.113** | **$\pm 0.201$** | **-0.15** | **$\pm 0.253$** |
| Ser | -0.086 | $\pm 0.079$ | -0.098 | $\pm 0.111$ | -0.074 | $\pm 0.069$ | -0.091 | $\pm 0.220$ |
| His | -0.059 | $\pm 0.160$ | 0.073 | $\pm 0.287$ | -0.042 | $\pm 0.104$ | -0.099 | $\pm 0.152$ |
| Gln | 0.179 | $\pm 0.484$ | 0.656 | $\pm 1.018$ | 0.274 | $\pm 0.646$ | 0.254 | $\pm 1.228$ |
| **Gly\*** | **-0.013** | **$\pm 0.109$** | **-0.032** | **$\pm 0.125$** | **0.036** | **$\pm 0.111$** | **0.102** | **$\pm 0.173$** |
| Thr | -0.016 | $\pm 0.088$ | -0.02 | $\pm 0.075$ | -0.033 | $\pm 0.105$ | 0.038 | $\pm 0.234$ |
| Arg | -0.729 | $\pm 0.556$ | -0.742 | $\pm 0.697$ | -0.669 | $\pm 0.449$ | -0.455 | $\pm 1.117$ |
| Ala | 0.865 | $\pm 0.492$ | 0.765 | $\pm 0.576$ | 0.748 | $\pm 0.650$ | 0.981 | $\pm 0.713$ |
| Tyr | -0.016 | $\pm 0.053$ | -0.034 | $\pm 0.056$ | -0.017 | $\pm 0.053$ | -0.009 | $\pm 0.071$ |
| Trp | -0.032 | $\pm 0.049$ | -0.026 | $\pm 0.057$ | -0.034 | $\pm 0.062$ | -0.061 | $\pm 0.075$ |
| Met | -0.031 | $\pm 0.027$ | -0.03 | $\pm 0.043$ | -0.018 | $\pm 0.024$ | -0.029 | $\pm 0.025$ |
| Val | -0.037 | $\pm 0.055$ | -0.039 | $\pm 0.072$ | -0.029 | $\pm 0.056$ | -0.032 | $\pm 0.059$ |
| Phe | -0.013 | $\pm 0.025$ | -0.012 | $\pm 0.042$ | -0.006 | $\pm 0.026$ | -0.011 | $\pm 0.029$ |
| Iso | -0.03 | $\pm 0.040$ | -0.028 | $\pm 0.06$ | -0.028 | $\pm 0.040$ | -0.034 | $\pm 0.047$ |
| **Leu\*** | **-0.086** | **$\pm 0.071$** | **-0.056** | **$\pm 0.128$** | **-0.027** | **$\pm 0.078$** | **-0.103** | **$\pm 0.086$** |
| Lys | 0.215 | $\pm 0.274$ | 0.383 | $\pm 0.462$ | 0.262 | $\pm 0.300$ | 0.261 | $\pm 0.276$ |

**Figure 3:**
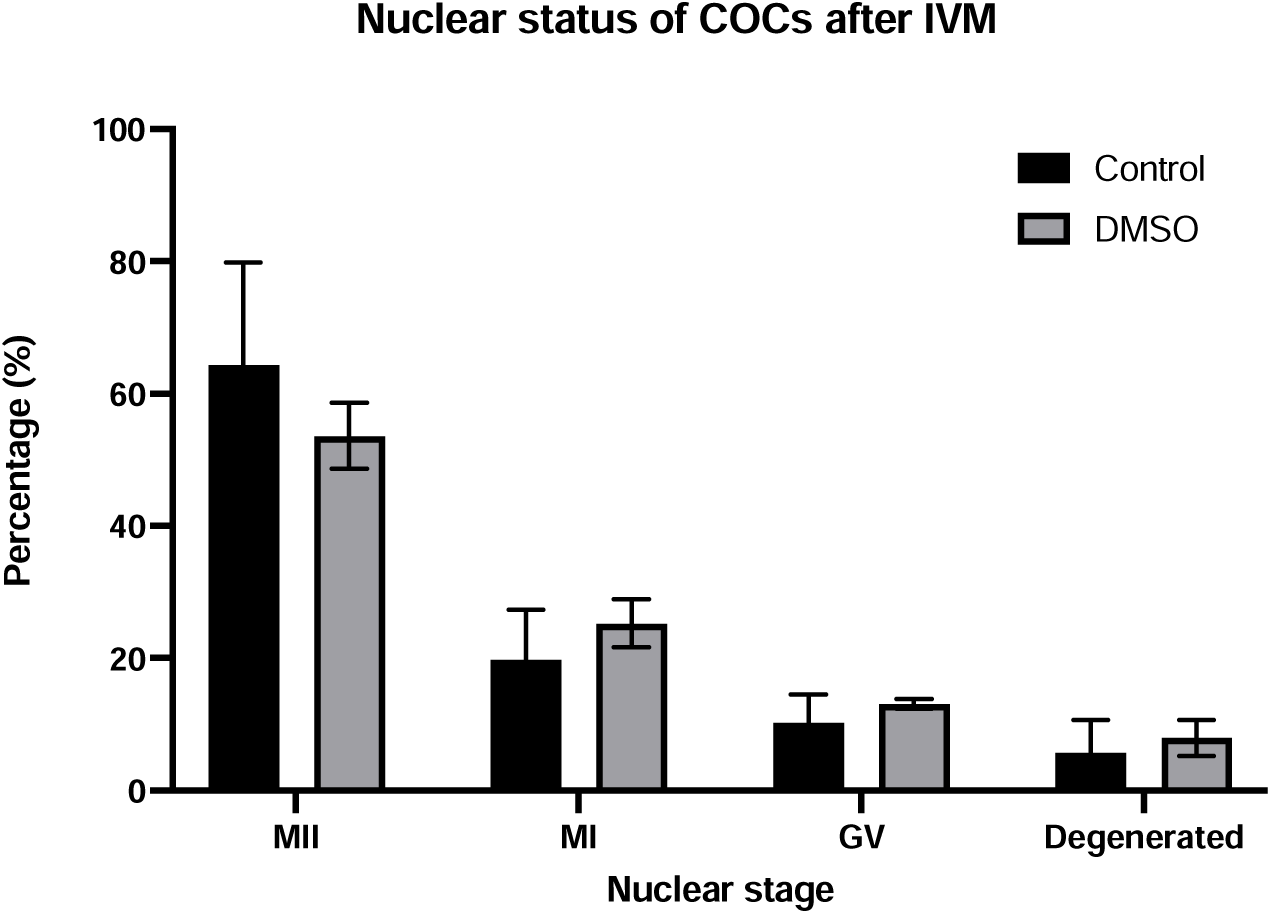
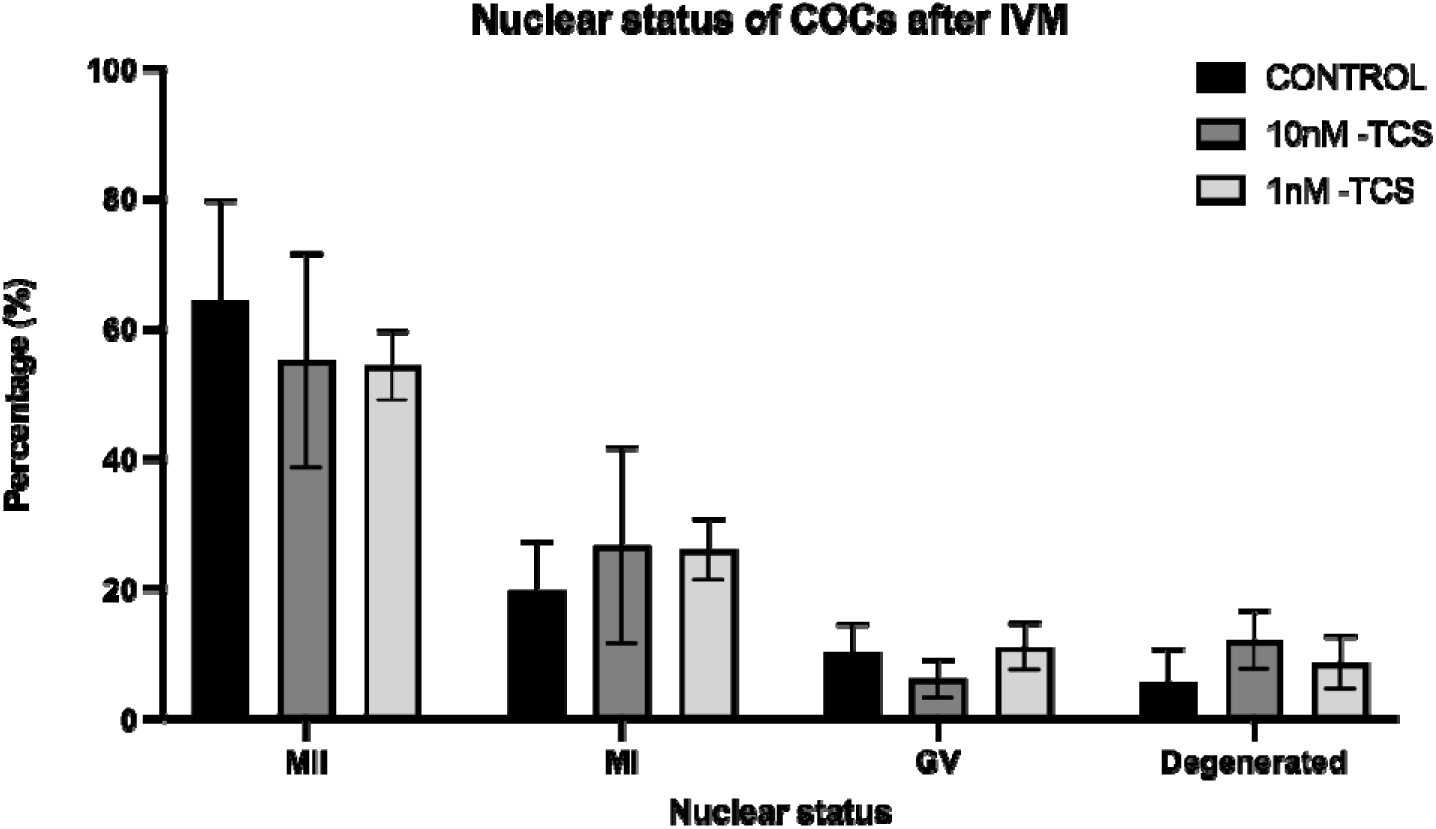
Nuclear status of bovine COCs exposed to TCS during IVM. Two-way ANOVA followed by Tukey test were performed. (n=3; 352 oocytes in total)

Since lactate production was not influenced, and the nuclear status of COCs after IVM was not significantly impacted by the presence of TCS, it was possible that the increased depletion of pyruvate by oocytes exposed to TCS exposure may reflect an increased demand for energy production in order to counterbalance the effects of the presence of TCS. In order to test this hypothesis, OCR and its component were measured on COCs during acute exposure to TCS prior to IVM.

The effect of TCS on bioenergetic profiles is presented in Figure 4. Basal OCR by COCs without TCS was first established, before TCS was injected. The OCR response to TCS was compared to basal respiration (Fig 4c). Subsequently, Oligomycin, as an ATP-synthase inhibitor, was injected into all groups allowing to indicate the proportion of OCR which is coupled to ATP generation. This was followed by the addition of FCCP which uncouples oxygen consumption from oxidative phosphorylation and thus indicates maximal OCR. Finally, Antimycin A, and Rotenone (A/R) were added in combination. The OCR insensitive to A/R is related to non-mitochondrial activity (Fig 4a). The response to the TCS treatment, and to the mitochondrial inhibitor per experimental group are illustrated in figure 4b.

**Figure 4:**
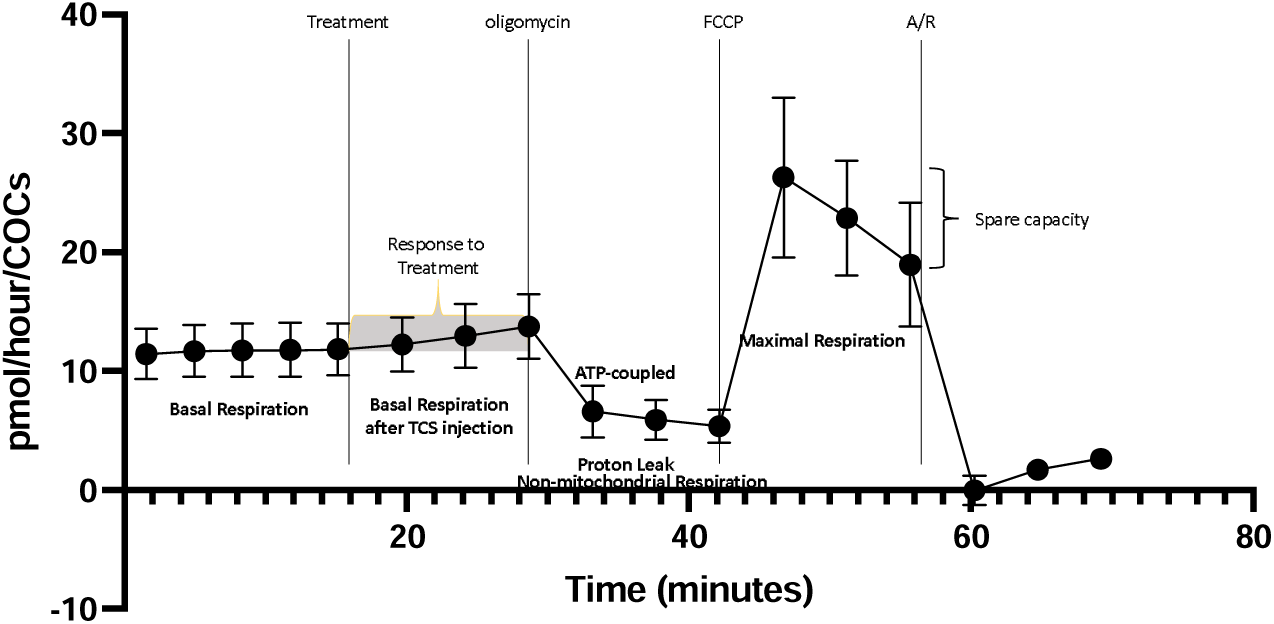
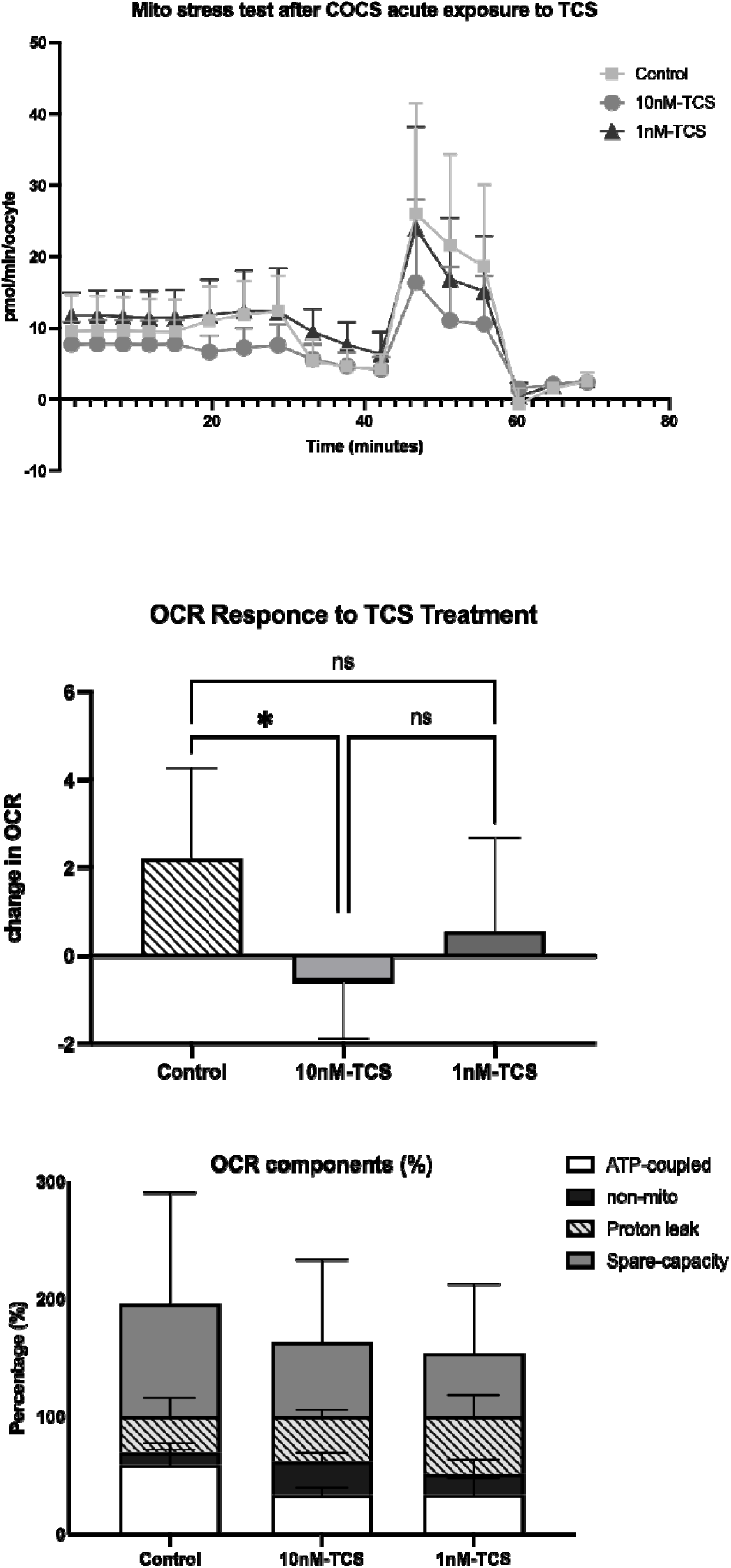
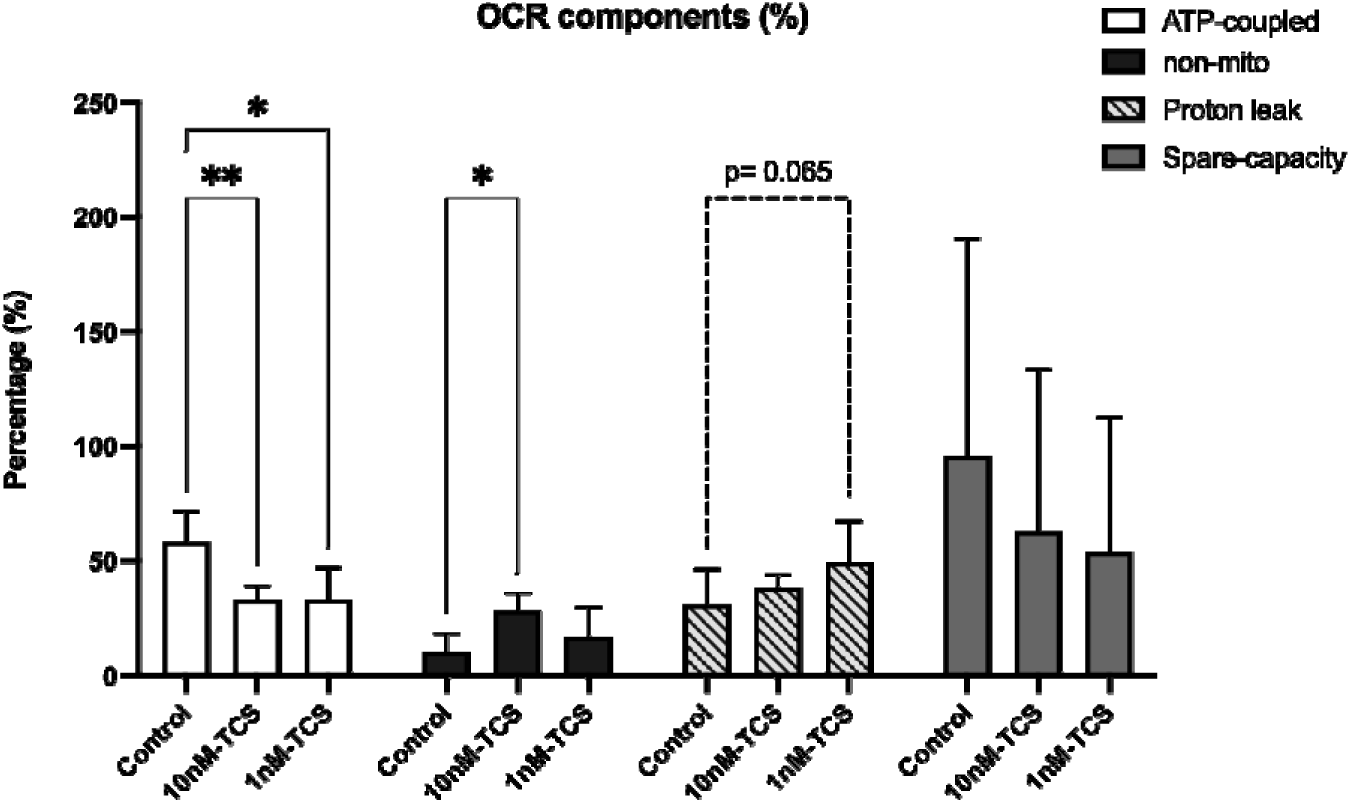
a) and b) OCR in COCs in response to TCS, oligomycin, FCCP and A/R a) as indicative examples of mitochondrial parameters and b) as comparison among the different treatments c) OCR as response to the TCS treatment, d) and e) mitochondrial parameters as a percentage of basal OCR after TCS injection. Differences were detected using c) one-way ANOVA followed by Tukey test. e) two-way ANOVA followed by Tukey test. * p<0.05; ** p<0.01; *** p<0.001; **** p<0.0001 (OCR, n=3).

**Figure 5:**
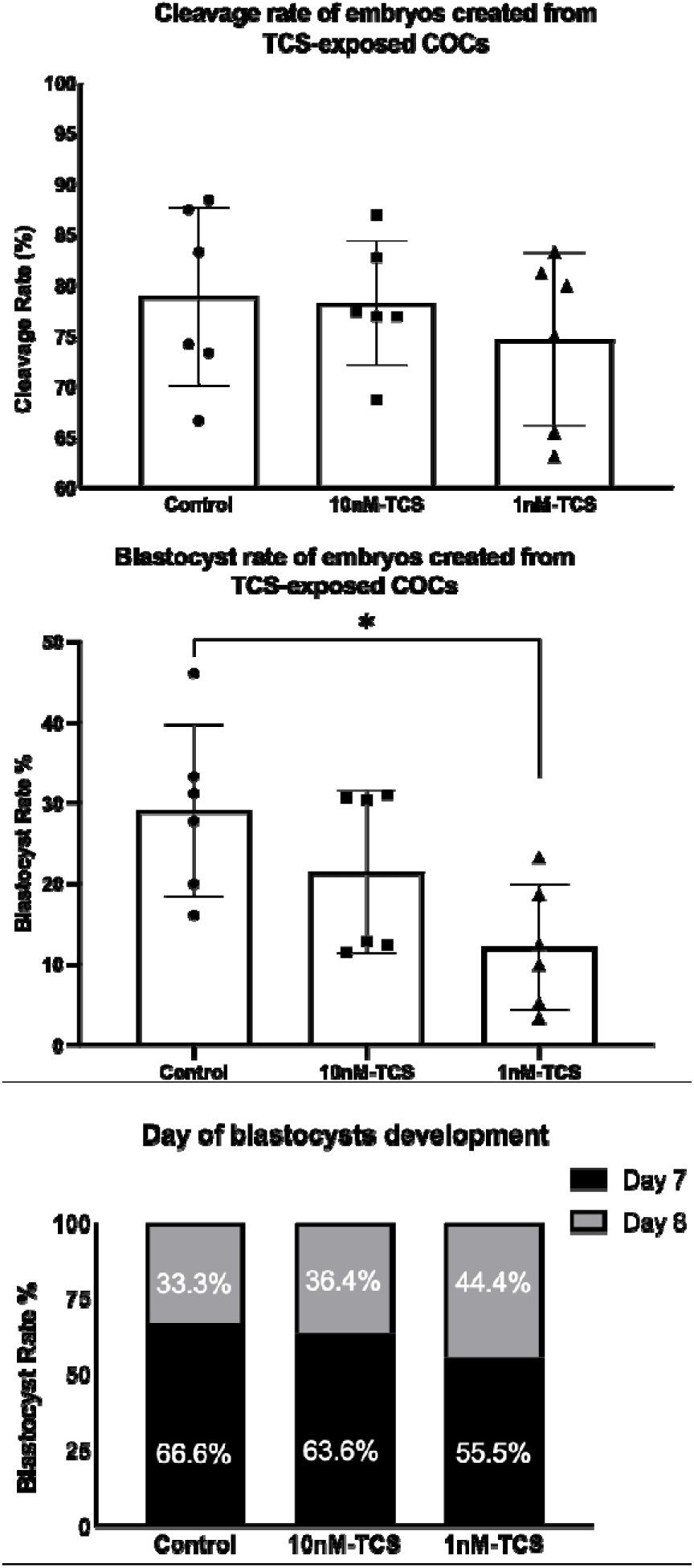
a) Cleavage rate and b) Blastocyst rate of embryos produced by TCS-exposed COCs during IVM. c) The day of blastocyst formation expressed as percentage of the total number of blastocysts per group. Differences were tested using one-way ANOVA and post hoc Tukey test. (n=6 biological replicates)

Upon addition of fresh culture medium to the control, there was a positive change (modest increase) in OCR, likely due to the replenishment of nutrients. However, this increase was ablated when 10 nM TCS was added, with a modest impairment of OCR (Fig 4c). By contrast, 1nM TCS had no effect on total OCR (Fig 4c).

The origin of the OCR difference in response to the treatment among the groups is presented in Figure 4d and 4e. The components of OCR were calculated as percentage to COCs basal respiration after the treatment’s injection. The proportion of OCR directly coupled to ATP synthesis was 58,7% ± 5.2% for control group whilst for 10nM-TCS and 1nM-TCS it was 33.1% ±2.5% and 33.3 ±5.6% respectively. In other words, oxygen consumption directly coupled to ATP synthesis was decreased by approximately 25% when the COCs were exposed to TCS irrespective of the administered dose; a statistically significant difference compared to the control group (Fig 4c). Moreover, the proportion of the OCR linked to non-mitochondrial activity was significantly increased when the COCs were exposed to 10nM-TCS dose (28.4% ± 3%) compared to the control group (10.5% ±3.2%). Finally, a strong trend of increased proton leak was observed when the COCs were exposed to the 1nM-TCS dose compared to control group (p=0.065).

TCS-treated COCs were next fertilised, and embryos cultured without further addition of TCS. Rates of cleavage and blastocyst formation were assessed on Day 2 and Day 7 to Day 8 respectively. The results showed that the cleavage rate of bovine embryos produced from TCS-exposed COCs was not significantly different to the control group. However, fewer blastocysts were formed from COCs-exposed to 1nM TCS compared to control group (12.22% vs 29.11% p=0.02). In addition to the reduced blastocyst rate in embryos derived from COCs-exposed to 1nM TCS, the observations hint/suggest? that the day of blastocyst formation was slightly delayed compared to control group, however this difference was not statistically significant. Future morphokinetic studies could possibly identify a more subtle difference in the timing of development.

Based on these findings, the metabolic profile of the blastocysts was also determined. To enable this, blastocysts were cultured in individual drops for 22 to 24 hours. The key cellular substrates, glucose, pyruvate and lactate, which are the main energy substrates during preimplantation embryo development, as well as 18 amino acids were all measured in spent media using a microfluorometric assay and HPLC, respectively.

Glucose consumption did not differ was significantly between blastocysts derived from TCS-exposed COCs compared to control group (Fig 6a). Further comparison of glucose depletion between blastocysts which developed and those which did not develop further during the 24 hour of individual culture showed, as was expected, that the developed blastocyst consumed higher amount of glucose per hour compared to those that did not develop. The difference of glucose consumption was statistically significant among the developed and non-developed blastocyst derived from the 10nM-TCS exposed COCs (Fig 6d).

**Figure 6:**
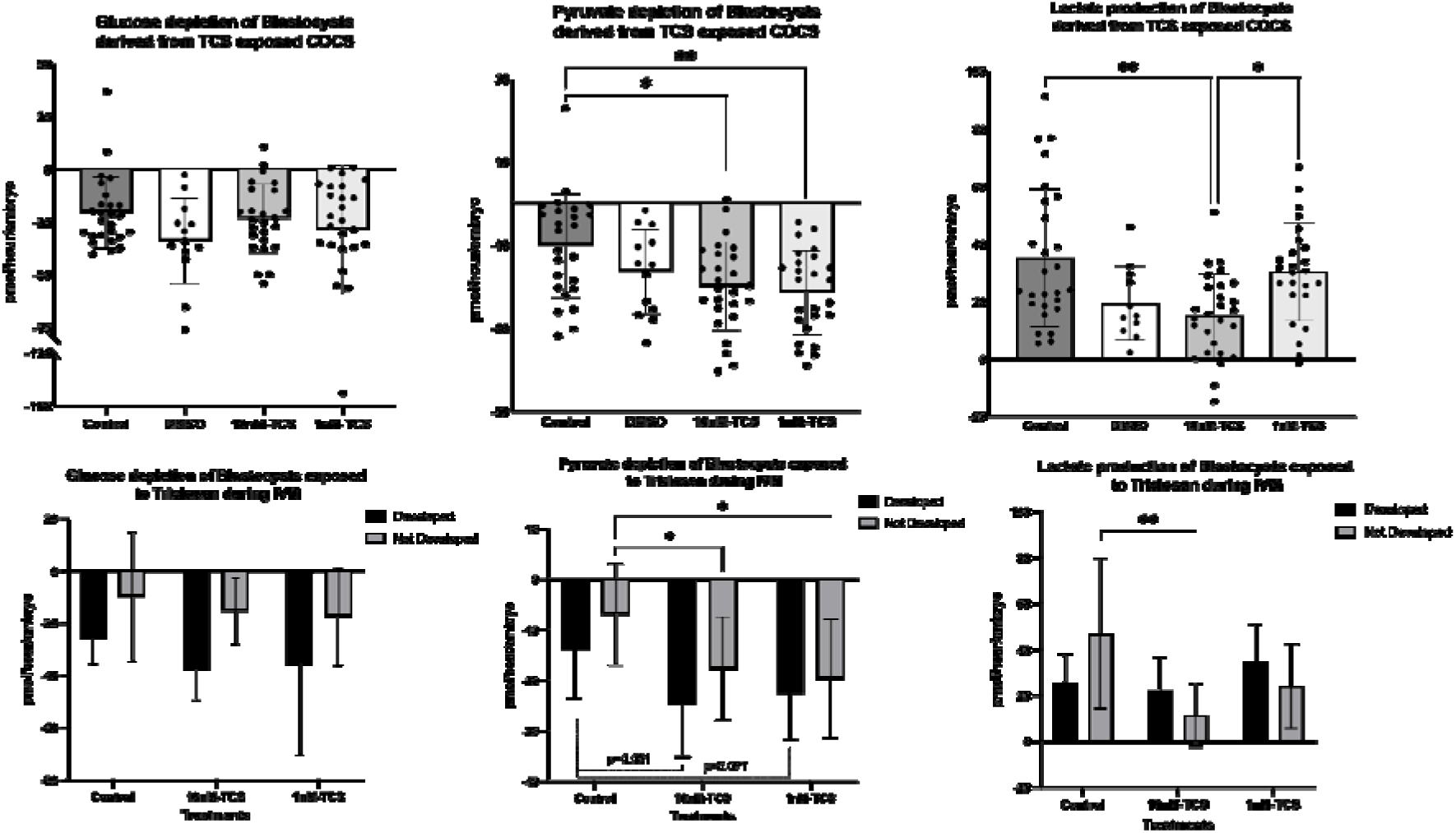
a) Glucose and b) Pyruvate depletion and c) Lactate production of the total number of blastocyst produced per treatment group. Each dot in graphs a), b), c) represents the energy substate’s production or depletion of an embryo. Comparison of d) Glucose e) Pyruvate depletion and f) lactate production by blastocysts whose development progressed (developed) or did not progress (not developed) during individual embryo culture. The distribution of each data set was tested; in those cases in which the distribution was nornal, a one-way ANOVA followed by Tukey test was perfomed (graph b) while the differernce between developed vs not developed was tested using two-way ANOVA (graph e). Kruskal-Wallis test followed by Dunn’s test was used when the data set failed in the normal distribution test (graphs a and c) In this case the differences among developed vs not developed blastocysts was tested using Kruskal-Wallis test followed by Dunn’s and Mann-Whitney (graph d and f) Data presented in pmol/hour/embryo ± SD.

However, the blastocysts derived from COCs treated with 10nM-TCS or 1nM-TCS consumed almost the double (1.96x) the amount of pyruvate compared to the blastocysts derived from the untreated COCs (−19.86 vs −10.12 pmol/hour/embryo p=0.011 and −21.41 vs −10.12 pmol/hour/embryo, respectively) (Fig 6b). Further analysis of these results shows that pyruvate production was significantly increased in the blastocyst group which remained in the same developmental stage during individual culture (not developed). (Fig 6e).

Finally, lactate production by blastocysts produced from the 10nM-TCS exposed COCs was significantly reduced compared to controls and to the low dose of TCS group (15.20 vs 31.82 pmol/hour/embryo and 15.20 vs 30.38 pmol/hour/embryo, respectively) (Fig 6c). However, the blastocysts with reduced lactate production had not progressed in their development during the preceding 24 hours of individual culture (Fig 6f)

Additionally, using the spent media after the 24 hours of individual blastocyst culture, amino acid turnover was determined by HPLC. The data revealed that, out of 18 amino acids measured, asparagine, glycine, leucine concentrations in spent medium of blastocysts from TCS-exposed COCs were significantly different to controls (Fig 7). Blastocysts derived from TCS-exposed COCs depleted asparagine and released glycine whilst the control groups released asparagine and consumed glycine. Interestingly, the low dose of TCS (1 nM) seemed to have a stronger effect on the metabolic state of those two amino acids compared to the higher dose (10 nM). Finally, leucine consumption was significantly decreased by blastocyst derived from COCs treated with the high dose of TCS compared to 1nM-TCS with a strong trend towards depletion compared to the control group.

**Figure 7:**
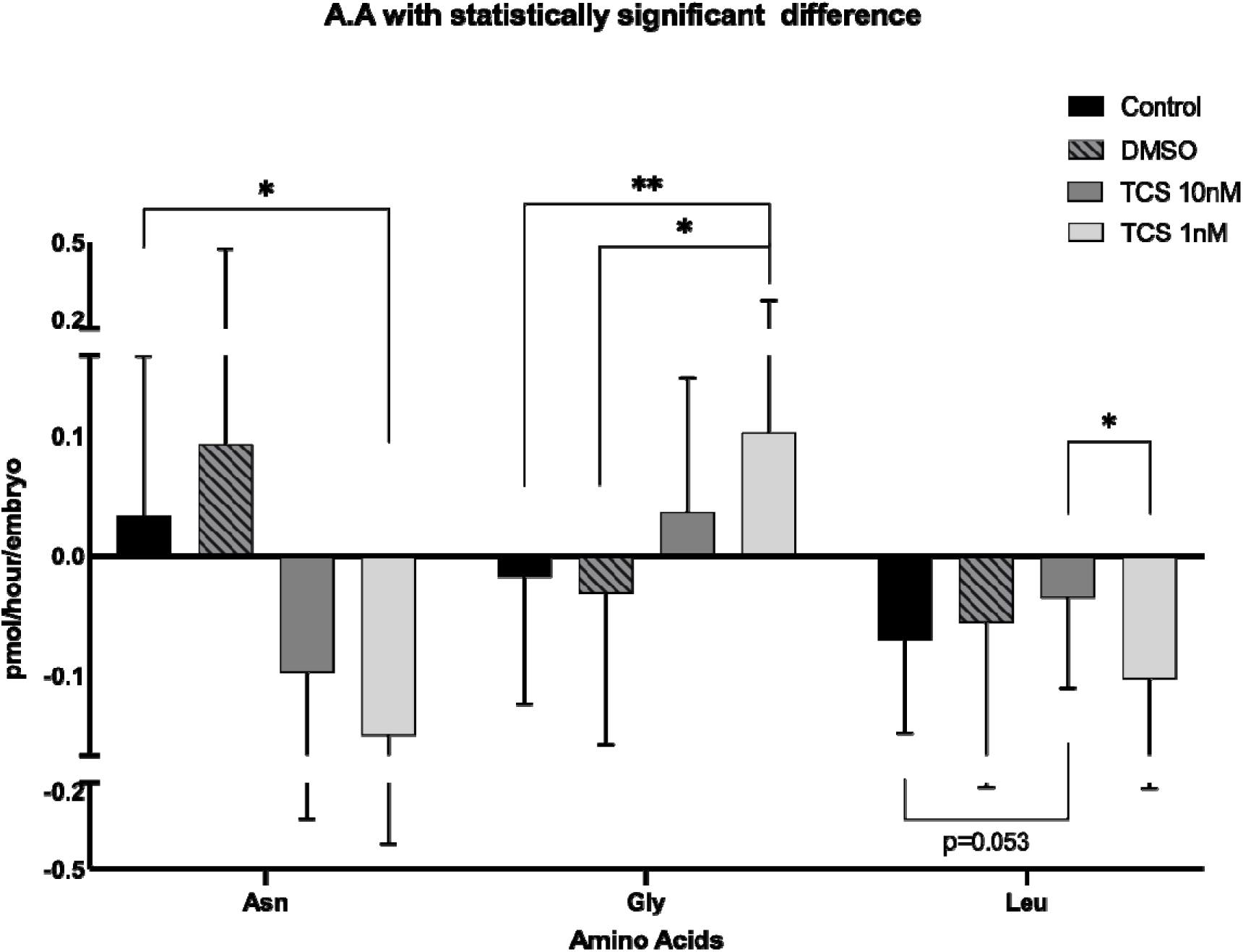
Amino acid turnover of blastocysts derived from TCS exposed COCs. The graph depicts the 3 out of 18 amino acids, Asparagine (Asn), Glycine (Gly),) and Leucine (Leu) with significant differences after statistical analysis using one-way ANOVA followed by post hoc analysis with Tukey or Kruskal-Wallis test followed by Dunn’s test depending on whether the amino acid followed a normal distribution or not. The results are expressed as pmol/hour/embryo ± SD.

## Discussion

In this study, we have used a bovine model to demonstrate that exposing oocytes to TCS leads to altered steroidogenesis and disrupted metabolic and mitochondrial function. In addition, embryos made from oocytes exposed to TCS exhibit significantly altered usage of metabolic substrates, indicative of further metabolic adaptation. The concentrations of TCS (1nM and 10nM) administrated are representative of TCS levels identified in human bodily fluids (Dayan, 2007; Allmyr *et al*., 2008; Calafat *et al*., 2008; Arbuckle *et al*., 2015; Azzouz *et al*., 2016, Li et al., 2025) which, in the majority of the cases, occur in the nanomolar range.

Oocyte quality and the environment in which development occurs are major determinants of ongoing embryo viability. The subsequent preimplantation stages of development are of critical importance since impairment during this period of development can have detrimental effects on pregnancy outcomes and, of greater concern, on lifelong health of the offspring, captured cogently by the Developmental Origins of Health and Disease hypothesis (Barker, 2007). According to this hypothesis, exposure of gametes and preimplantation embryos to chemicals and other environmental agents can alter the developmental fate of an organism.

Due to industrialization and modern lifestyle changes, humans and other organisms, are exposed to diverse classes of chemicals either indirectly due to their accumulation in the environment (Zhang *et al*., 2021) and food (Darnerud *et al*., 2006; Rashid *et al*., 2020) or directly through use of consumer products (Dionisio *et al*., 2018). A typical example of this is Triclosan (TCS), a chemical which has been used in many personal hygiene products as an antimicrobial agent (Weatherly and Gosse, 2017) and may accumulate in the environment due to its excessive use (Bedoux *et al*., 2012; Weatherly and Gosse, 2017). Thus, as mentioned earlier, TCS has been identified in human biological fluids such as urine, serum and breast milk and ovarian follicular fluid (Li *et al.,* 2024).

In addition to the results presented herein, a spectrum of *in vitro* and *in vivo* studies using TCS exposures have shown detrimental effects on different physiological endpoints (Weatherly and Gosse, 2017; Shrestha *et al*., 2020), including the reproductive system (Zhu *et al*., 2016, 2019). Specifically, such studies have revealed that TCS acts as Endocrine-Disrupting Chemical (EDC) and can interfere with E2, P, and T4 production due its similarity with 17β-estradiol (Wang and Tian, 2015). Despite these findings, the impact of TCS exposure on mammalian oocyte maturation and the early stages of embryo development remain largely unknown. Two of very few papers in this area (Park *et al*., 2020; Zhao *et al*., 2025) examined TCS toxicity in porcine oocytes and reported results which aligned with our own findings in suggesting that TCS exposure during in vitro maturation impairs mitochondrial function in porcine oocytes. It has also been reported that preimplantation embryo development in pigs (Kim *et al*., 2020) and mouse (Yang *et al*., 2022) is influenced by the presence of TCS. Despite growing evidence that TCS has negative impact in early reproductive events, due to the use of a wide range administrated concentrations and periods of exposure that have been used, the precise mechanism of action of TCS remains unclear.

The results of the present study revealed that bovine COCs treated with 1nM TCS released significantly more progesterone than controls. Similar observations have been reported previously for human (Du *et al*., 2021) and rat (Chen *et al*., 2019) granulosa cells in a range of different TCS concentration (1 nM −10 μM)). However, when rats were supplemented with TCS in food at 50 mg/kg/day for 28 consecutive days, progesterone levels in serum fell significantly (Arismendi *et al*., 2022). Progesterone levels decrease have been reported in mice in vivo exposed to 10 and 100mg/kg/day (Cao *et al*., 2018).

Moreover, exposure to TCS (1 nM −10 μM) has been reported to increase production of E2 in human and rat granulosa cells (Chen *et al*., 2019; Du *et al*., 2021). This contrasts our observations; 17β-estradiol production during in vitro maturation of bovine COCs remained unaffected by TCS. However, our results do align with findings from whole animal dose studies in rat (Arismendi *et al*., 2022)and mouse (Cao *et al*., 2018), who failed to see any differences in E2 levels in serum in response to TCS.

The apparent differences observed in our study providing evidence for an influence of TCS on P4 and E2 levels may be explained by the duration of the exposure or the animal model that have been used. In addition, , this/these differences?? may arise from different administered doses, indicating a nonmonotonic action of TCS. The nonmonotonic action of TCS can further be supported from our results as the 1nM dose affected P4 production however this remained unchanged when the COCs were exposed to 10nM TCS. The nonmonotonic action of EDCs has previously been reported (Vandenberg *et al*., 2012), notably for BPA, (Vandenberg, 2014). TCS has a similar chemical structure to BPA, both of which exhibit estrogenic action and structural similarity to 17β estradiol. Importantly, our findings align with the conclusion of studies suggesting the endocrine disrupting properties of TCS. However, further studies on the non-monotonic action of TCS need to be done.

Beyond steroidogenesis, exposure to TCS significantly affected the metabolic status of COCs in culture. COCs exposed to 1nM or 10nM TCS after IVM depleted pyruvate from the culture medium, whereas controls released pyruvate. Pyruvate is an essential energy substrate during preimplantation embryo development (Nagaraj *et al*., 2017). Furthermore, pyruvate is required for the completion of meiotic maturation in mouse (Downs *et al*., 2002; Johnson *et al*., 2007). In a study conducted in mouse COCs using maturation media supplemented with FSH during IVM, pyruvate was produced by COCs that had reached the MII stage but was consumed by nuclear immature COCs (Roberts *et al*., 2004). Pyruvate is also produced as the product of glycolysis in the cytoplasm from where it is either converted to lactate or enters the mitochondria and is converted to acetyl-CoA which enters the TCA cycle. Based on our observations of altered pyruvate handling by COCs treated with TCS, we proposed that these COCs were either entering meiotic arrest or displaying mitochondrial dysfunction. To exclude meiotic arrest as the cause for disrupted pyruvate metabolism, the nuclear status of COCs exposed to TCS was determined. The results confirmed that maturation status was not influenced by TCS treatment. We therefore examined mitochondrial function by determining Oxygen Consumption Rate (OCR). Immediately after aspiration, COCs were treated with 0, 1 or 10nM of TCS, and OCR determined. In addition, mitochondrial function was examined using the mitochondrial inhibitors oligomycin and Antimycin/Rotenone mix which inhibit Complex V, and Complex III/Complex I respectively, as well as FCCP which acts as a mitochondrial uncoupler (Muller *et al*., 2019). The results revealed that the total OCR was significantly reduced in COCs exposed to 10nM dose of TCS. Further analysis of this result showed that the ATP-coupled OCR when expressed as a percentage of the total OCR was significantly reduced in TCS-treated COCs compared to the untreated group. Mitochondrial dysfunction due to decreased ATP levels after TCS exposure have also been reported recently in porcine oocytes (Zhao *et al*., 2025). In addition, the rate of non-mitochondrial OCR was significantly increased after 10nM TCS exposure and a strong increase observed in the proportion of OCR devoted to proton leak after COCs exposure to 1nM TCS. These results confirm the mitochondrial dysfunction after TCS exposure of bovine COCs during IVM. TCS mitochondrial toxicity has previously been confirmed in different mammalian cells including porcine sperm (using exposure levels of 1-10μg/ml; (Ajao *et al*., 2015)), human granulosa cells (Du *et al*., 2021) and porcine oocytes and embryos (Kim *et al*., 2020; Park *et al*., 2020) after direct exposure in the micromolar range (from 1 to 100μΜ).

To investigate if onward oocyte developmental competence is impaired aftereTCS exposure, the impact on the early stages of embryo development of TCS-exposed COCs was investigated. COCs treated with TCS were fertilised and cultured for seven days without further exposure to TCS. Our findings revealed that blastocyst rate was significantly reduced in the 1nM-TCS group compared to control group (12.22% vs 29.11% p=0.0205). However, a dose of 10nM-TCS did not influence the developmental rates of the embryos. Reduced blastocyst rate due to TCS exposure during preimplantation stages of development has recently been confirmed in a study using the porcine as a model (Reference?. In this? study, mitochondrial dysfunction was associated with a low blastocyst rate which was partially confirmed by our results where we observed that both doses of TCS impaired mitochondrial function, but that the 1nM dose of TCS only had an impact on blastocyst rate.

Finally, the metabolic status of the blastocysts generated in each group was assessed by transferring them in individual drop culture system for 24 hours. The blastocysts derived from TCS-treated COCs consumed significantly more pyruvate, irrespective of the administrated dose compared to controls group. Meanwhile, the lactate production of blastocysts derived from 10nM-TCS COCs was significantly reduced. Further analysis showed that pyruvate depletion was higher compared to the control group, regardless of the development of blastocyst during the 24-hours of individual culture (? don’t get this). Finally, the amino acid turnover of the blastocysts generated was influenced in that 3 out of 18 amino acids were significantly different among the groups.

Overall, our results show that the metabolic activity of blastocysts derived from TCS treatment were in a “hyperactivated mode” in that they consumed more pyruvate and increased the turnover of the amino acids. One interpretation of these data is that embryos derived from COCs treated with TCS increase their metabolic function in order to counterbalance the effects of TCS. Embryos with higher metabolic activity in terms of pyruvate consumption (Guerif *et al*., 2013) amino acids turnover (Houghton *et al*., 2002; Brison *et al*., 2004) and oxygen consumption (Lopes *et al*., 2007, 2010) have previously been associated with lower developmental potential (Leese *et al*., 2022)

Human females with subfertility may use Assisted Reproductive Techniques (ART) however, the etiology of about 15% of the infertility cases remain unclear (Comhaire, 1987). Subfertility or infertility of human females at reproductive age has been correlated with a wide range of causes/factors? including exposure to natural or synthetic chemicals (Pizzorno, 2018). The bovine has previously been suggested as a suitable model for human IVF (Ménézo and Hérubel, 2002) and to evaluate the effect of chemicals on toxicological studies in reproduction (Santos *et al*., 2014) due to the biochemical and developmental similarities during the final stages of oocyte maturation, fertilisation and early stages of embryo development. As mentioned earlier, TCS, due to its excessive use, is accumulated in the environment resulting on people receiving direct or indirect exposure. For this reason, the present study was conducted to test TCS effects on oocyte and embryo metabolism using the bovine as a model for human.

Overall, this study has found that TCS exposure during IVM impairs the metabolic status of COCs by affecting the progesterone production and mitochondrial function resulting in reduced developmental competence. The COCs’ altered metabolic function not only reduced the developmental rates but also interfered with the metabolic activity of the embryos produced resulting in reduced developmental potential. These findings are of special concern taking into consideration that the levels of TCS chosen for the present study reflect levels common in everyday exposures for humans. Crucially, the implications of these findings in the longer term are not known but may relate to adverse effects on pregnancy outcomes or on health in later stage of development. Future studies could examine the action of TCS and related compounds on embryo morphokinetics and metabolism during the early stages of embryo development and during the onset of implantation and the impact of this exposure on later stages of life.

## Acknowledgment

This work was funded by the University of Hull. In addition, we would like to thank ABP York for providing the bovine reproductive tracts for these experiments

## Author contributions

V.P. designed experiments, performed experiments, analysed data, wrote manuscript. P.MK designed experiments, J.M.R designed experiments. H.J.L designed the experiments. R.G.S. Conceived study, designed experiments, wrote manuscript. All authors commented on manuscript during preparation.

## Competing interests

The authors declare no competing interests.

## Notes

### Competing Interest Statement

The authors have declared no competing interest.

